# Na^+^-translocating oxaloacetate decarboxylase from *Vibrio cholerae*: the functional tautomeric form of the substrate and the proton pathways in catalysis

**DOI:** 10.64898/2026.06.08.730933

**Authors:** Yulia V. Bertsova, Alexander D. Kvartalov, Marina V. Serebryakova, Alexander A. Baykov, Alexander V. Bogachev

**Affiliations:** Belozersky Institute of Physico-Chemical Biology, Lomonosov Moscow State University, 119234 Moscow, Russia

**Keywords:** sodium pump, proton transport, keto-enol tautomerization, general acid catalysis, oxalate, pyranine

## Abstract

Membrane-bound decarboxylases couple carboxylic acid decarboxylation to the transport of Na^+^ ions out of prokaryotic cells. The molecular mechanism of decarboxylase action is not yet known, which contrasts with the progress achieved in studying other primary ion pumps. Measuring decarboxylase activity is complicated by slow keto-enol tautomerization of the substrates during the assay. We found that HEPES exhibits anomalously high efficiency as a general acid catalyst for C-H bond formation during the enol-to-ketone conversion of oxaloacetate. Accordingly, the addition of HEPES to the assay medium eliminated the contribution of tautomerization rate to measured decarboxylation rate. Using the dependence of oxaloacetate tautomerization rate and equilibrium on solvent properties and pH, we established that only the keto form of oxaloacetate is converted by *Vibrio cholerae* oxaloacetate decarboxylase. Steady-state kinetic measurements did not reveal cooperativity in oxaloacetate conversion and Na^+^ binding. The effects of ionophores (CCCP, valinomycin, and ETH157) on proton transport in pyranine-loaded membrane vesicles prepared from *V. cholerae* cells indicated that the proton required for the conversion of oxaloacetate to pyruvate is taken up from the cytoplasmic side of the membrane. Furthermore, the effects suggested that ΔpH generation is caused by secondary electrophoretic proton transport in exchange for Na^+^.These findings advance our understanding of the molecular mechanism of the decarboxylation-supported Na^+^ transport in bacteria.

## 1. INTRODUCTION

The decarboxylation of carboxylic acids is energetically favorable (ΔG^°^’ = –(20 – 30) kJ/mol) [1, 2]; however, the released energy is insufficient for ATP synthesis via substrate-level phosphorylation. Unique membrane-bound Na^+^-translocating decarboxylases convert this energy into a transmembrane Na^+^ electrochemical potential difference [2]. The latter can be further used for ATP synthesis, the transport of substances across the membrane, or bacterial flagellar rotation [3]. To date, four types of such decarboxylases have been described, which use oxaloacetate, malonate, methylmalonyl-CoA, or glutaconyl-CoA as the substrate [1, 4]. The best-known of these is Na^+^-translocating oxaloacetate decarboxylase (OAD, EC 7.2.4.2), which participates in the anaerobic fermentation of unusual substrates, such as citrate or tartrate, in many bacteria [4].

Historically, the identification of OAD as the first sodium pump [5] led to the discovery of the sodium cycle of energy conversion [3]. This enzyme, found in many bacteria and archaea, appears to be a relic of the bioenergetic and metabolic pathways of the early forms of life [6]. OAD is important for the pathogenicity of some contemporary disease-causing bacteria [7, 8] and may be a promising molecular target in biomedicine.

The OAD molecule is a trimer or tetramer of trimers [9, 10], each formed by the α (63 kDa), β (45 kDa), and γ (9 kDa) subunits [11]. The water-soluble α subunit is exposed to the cytoplasm and consists of three domains (carboxyltransferase, association, and biotin-binding) connected by flexible linkers rich in proline and alanine residues [12]. This subunit contains biotin and 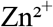 as prosthetic groups and catalyzes oxaloacetate decarboxylation to yield pyruvate and carboxybiotin, the latter being covalently attached to the biotin-binding domain of the α subunit. The β subunit is located in the cytoplasmic membrane and contains 10 transmembrane α-helices and 2 intramembrane helical hairpins, forming an inverted topological repeat [10], a feature characteristic of membrane transporters using an alternating access mechanism. This subunit is involved in the decarboxylation of carboxybiotin and carries out the coupled transmembrane transfer of sodium ions. The γ subunit contains one transmembrane α-helix and links the α and β subunits together [10, 13].

OAD catalyzes a two-step reaction of oxaloacetate decarboxylation (Fig. 1), coupled to the transfer of two Na^+^ ions across the membrane [14]. The first step involves carboxyl group transfer from oxaloacetate to biotin in the carboxyltransferase domain of the α subunit. Subsequently, N-carboxybiotin is decarboxylated in a reaction involving the β subunit. The first step does not require Na^+^ whereas the second step is Na^+^-dependent and, hence, coupled to Na^+^ transport [15]. The molecular mechanisms of the chemical and transport reactions are still poorly understood. Oxaloacetate exists in two tautomeric forms (ketone and enol) (Fig. 1), and it remains to be determined which of them is the true substrate. The decarboxylation reaction consumes a proton, which is currently believed to be taken up from the extracellular medium [16], a view that lacks strong experimental support.

**Fig. 1.**
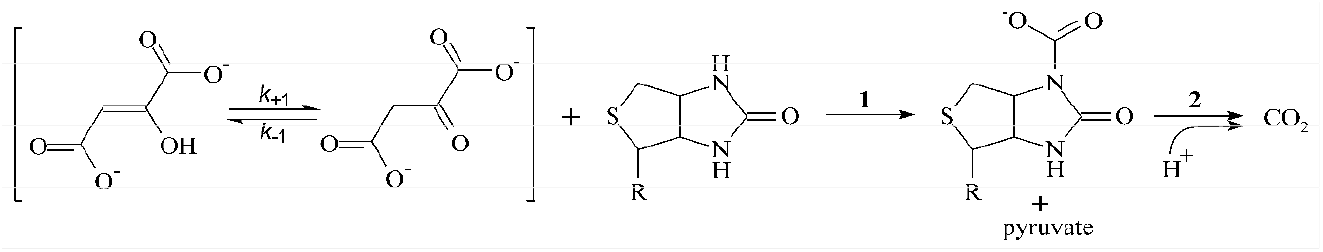
Scheme of the two-step reaction of oxaloacetate decarboxylation catalyzed by OAD.

The genome of the marine pathogenic bacterium *Vibrio cholerae* contains two *oad* operons encoding membrane oxaloacetate decarboxylases [9]. The physiological function of the *oad-1* operon product and the conditions for its induction are unknown. The *oad-2* operon is located within a cluster of genes required for citrate fermentation. Its product (OAD-2) is accumulated in large amounts during anaerobic growth of *V. cholerae* on citrate, resulting in very high oxaloacetate decarboxylase activity (> 1 U/mg protein) in the membrane fraction of the cells [9]. This feature and the stability of OAD-2 make it a convenient model for studying oxaloacetate decarboxylases [9].

The purpose of this investigation was to re-examine the OAD assay and use it to identify the tautomeric form of the substrate recognized and converted by OAD-2. Furthermore, we used membrane ionophores to determine the origin of the “catalytic” proton. The findings seem critically important for our understanding of the mechanism of the decarboxylation-supported Na^+^ transport in bacteria.

## 2. MATERIALS AND METHODS

### 2.1. Bacterial strains and growth conditions

The non-pathogenic *V. cholerae* strain O395N1 Δ*toxR* Δ*tcpP* [17] was used in this work. To induce OAD-2 synthesis, *V. cholerae* cells were grown anaerobically using citrate as the sole fermentable substrate [9]. Cultivation was carried out at 32 °C until mid-logarithmic growth phase in a medium containing 15 g/L NaCl, 0.75 g/L KCl, 1.2 g/L MgSO_4_·7H_2_O, 1 g/L (NH_4_)_2_SO_4_, 50 mM Tris-HCl (pH 8.0), 30 mM sodium citrate, 0.1 mM KH_2_PO_4_, 0.05% yeast extract, and 50 µg/mL streptomycin.

### 2.2. Isolation of membrane vesicles

Harvested *V. cholerae* cells were pelleted by centrifugation (7,500 *g*, 10 min) and washed with a medium containing 300 mM NaCl, 10 mM Tris-HCl (pH 8.0), and 5 mM MgSO_4_. The cell pellet was then resuspended in a medium containing 250 mM NaCl, 50 mM Tris-HCl (pH 7.5), 5 mM MgSO_4_, and a small amount of DNase. The cell suspension was passed through a French press (16,000 p.s.i.). Unbroken cells were pelleted at 20,400 *g* for 5 min. Membrane vesicles were sedimented by centrifugation at 140,000 *g* (60 min), the pellet was resuspended in a medium containing 100 mM KCl, 50 mM Tris-HCl (pH 7.5), and 5 mM MgSO_4_, frozen in liquid nitrogen, and stored at –70 °C. Protein concentration was determined by the bicinchoninic acid method [18] using bovine serum albumin as a standard.

To obtain membrane vesicles with encapsulated pyranine, washed *V. cholerae* cells were resuspended in buffer A (100 mM MOPS-Tris (pH 6.5), 25 mM K_2_SO_4_, 5 mM MgSO_4_) containing 2 mM pyranine (Eastman Kodak, USA) and a small amount of DNase. The cells were disrupted using a French press (16,000 p.s.i.). Unbroken cells were pelleted at 20,400 *g* for 5 min. Membrane vesicles were obtained by centrifugation at 140,000 *g* (60 min). The vesicles were then washed twice with buffer A by resuspension and pelleting at 140,000 *g* (50 min). The final pellet was resuspended in buffer A and used for ΔpH generation measurements on the day of preparation.

### 2.3. Purification of OAD-2 from the membrane vesicles

Purification of OAD-2 was performed by affinity chromatography on avidin-agarose using a previously described method [19] with minor modifications. Tetrameric avidin coupled to agarose (Imtek, Russia) was converted to the monomeric form by treatment with guanidine hydrochloride [20]. A suspension of membrane vesicles (15 mg protein/mL) in buffer (250 mM NaCl, 50 mM Tris-HCl (pH 7.5)) was solubilized with 2% Triton X-100 (Sigma-Aldrich Co, USA) and centrifuged at 140,000 *g* (40 min). The supernatant was applied to a column containing 2.5 mL of the affinity matrix equilibrated with buffer B (100 mM NaCl, 50 mM Tris-HCl (pH 7.5), 0.03% Triton X-100). The column was washed with 8 mL of buffer B, and OAD-2 was eluted with buffer B containing 2.5 mM D-biotin (CDH, India).

SDS-PAGE of OAD-2 was performed using 12.5% polyacrylamide gels [21]. Gels were stained for protein with PageBlue™ solution (Fermentas, Lithuania). MALDI-TOF MS analysis of protein bands was carried out on an UltrafleXtreme MALDI-TOF-TOF mass spectrometer (Bruker Daltonics, Germany) as described previously [22].

### 2.4. Determination of OAD-2 activity

OAD-2 activity in membrane vesicles was determined by measuring oxaloacetate consumption [23, 24] in dual-wavelength mode (260 – 290 nm, ε = 1.13 mM^−1^ cm^−1^) using an Aminco DW-2000 spectrophotometer (USA) at 25 °C. The assay medium contained 100 mM HEPES-Tris (pH 7.5), 0.08 – 50 mM NaCl, and 5–350 μM oxaloacetate (Sigma-Aldrich Co). The background sodium concentration in the media used was determined by flame photometry.

The activity of purified OAD-2 was determined at 25 °C in a coupled assay with lactate dehydrogenase (LDH, Sigma-Aldrich Co) by measuring the decrease in NADH absorbance (ε_340_ = 6.2 mM^−1^ cm^−1^) [23, 24] using a Hitachi-557 spectrophotometer (Japan). The assay medium contained 20 mM NaCl, 1 U/mL LDH, 0.1 mM NADH, 50 μM oxaloacetate, and either 100 mM HEPES-KOH or 2 mM Tris-HCl. Special care was taken to maintain pH 7.5 after the addition of oxaloacetic acid in both cases.

### 2.5. Generation of ΔpH in V. cholerae membrane vesicles

Alkalization of the vesicle interior was monitored by measuring the change in encapsulated pyranine fluorescence [25] at 510 nm (excitation at 458 nm). Measurements were carried out in a medium containing 100 mM MOPS-Tris (pH 6.5), 25 mM K_2_SO_4_, 5 mM MgSO_4_, membrane vesicles with entrapped pyranine (75 μg protein per mL) in the presence or absence of 10 mM Na_2_SO_4_. After a 5-min preincubation of the vesicles, the reaction was initiated by the addition of 0.3 mM oxaloacetate. Fluorescence was recorded at 25 °C using a FluoroMax-3 spectrofluorometer (Horiba Jobin Yvon, France).

## 3. RESULTS AND DISCUSSION

### 3.1. Optimization of the method for measuring OAD activity

Two methods for measuring OAD activity have been proposed. In one of them, the reaction product, pyruvate, is monitored by measuring NADH consumption in a coupled assay with lactate dehydrogenase (LDH) [23, 24]. This method is sensitive but applicable only to purified OAD preparations because of the presence of interfering malate dehydrogenase activity in crude preparations, such as bacterial membranes. In membrane preparations, OAD activity is determined by measuring the absorbance of the substrate oxaloacetate at around 260 nm [23, 24]. This wavelength corresponds to the absorption maximum of the enol form of this compound (the keto form of oxaloacetate absorbs light at a considerably shorter wavelength). This method is characterized by lower sensitivity. Importantly, the use of both methods is complicated by the low rate of keto-enol tautomerization of oxaloacetate [26], which can distort the measured kinetics of OAD activity, especially when assaying high OAD activities [23]. We therefore attempted to optimize the OAD assay by varying the nature and concentration of the pH buffer.

The tautomerization equilibrium of oxaloacetate depends on the solvent used and, to a lesser extent, on the pH value. Thus, a solution of oxaloacetic acid in dry diethyl ether contains 67% of the enol form, which drops to 13% in an aqueous medium at neutral and alkaline pH values and to 6.5% at pH 2 [27]. Hence, the transfer of oxaloacetic acid from diethyl ether or from an aqueous solution with a low pH value to an aqueous solution with a near-neutral pH (Fig. 2A and 2B, respectively) causes a shift in the equilibrium position and allows the kinetics of keto-enol tautomerization to be measured in both directions [27].

**Fig. 2.**
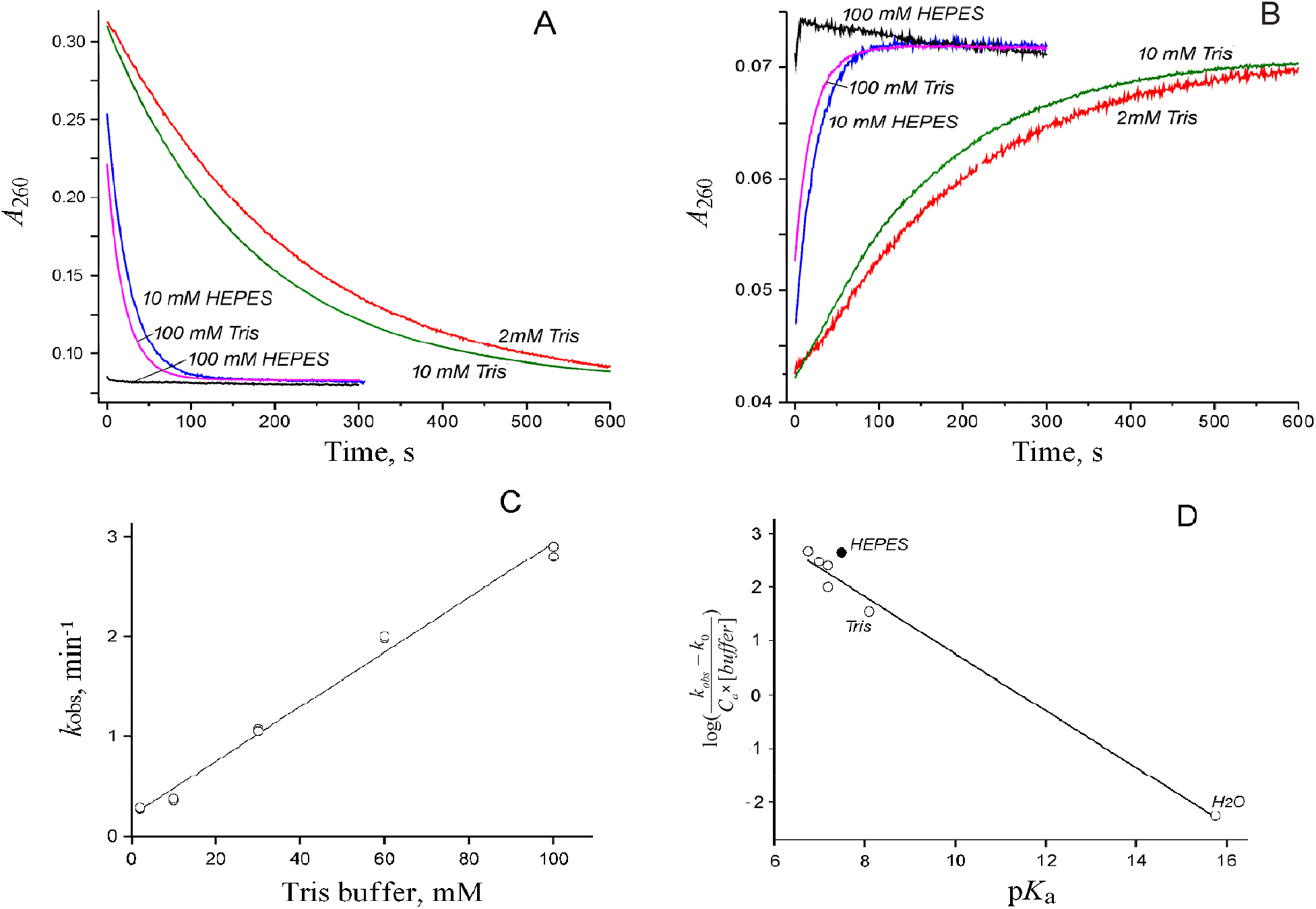
Kinetics of the non-enzymatic keto-enol tautomerization of oxaloacetate, measured by following changes in enol content at pH 7.5. **(A)** Kinetics of the tautomerization in the direction enol→ketone. Oxaloacetic acid (40 μM) was added from a 50 mM stock solution prepared in dry diethyl ether to a medium containing Tris-HCl or HEPES-KOH at the indicated concentrations. **(B)** Kinetics of the reaction in the direction ketone→enol. The experimental conditions were as for panel A, but oxaloacetic acid was added from an aqueous stock solution maintained at pH 2. **(C)** Effect of Tris buffer concentration on the apparent rate constant of oxaloacetate tautomerization at pH 7.5. **(D)** The Brönsted plot for the general acid catalysis of oxaloacetate tautomerization, constructed from the data in Table 1. The point for water was obtained using the *k*_obs_ value extrapolated to zero Tris concentration in Fig. 2C, the water concentration of 55.5 M, and p*K*_a_ of 15.7 for the H_2_O ↔ OH^−^ + H^+^ equilibrium. The straight line was drawn by the least-squares method using all points, except for the HEPES point (filled circle).

The tautomerization rate was characterized in terms of the apparent first-order rate constant *k*_obs_, obtained by fitting the rate equation for a reversible first-order reaction:

**Table 1.** Apparent rate constants of oxaloacetate tautomerization in the presence of various pH buffers at pH 7.5.

| Buffer | $pK_a^a$ | $C_a^b$ | $k_{obs} = k_{+1} + k_{-1}^c, \text{ min}^{-1}$ | | $(k_{obs} - k_0)/(C_a \times [\text{buffer}])^d, \text{ M}^{-1} \text{ min}^{-1}$ |
| --- | --- | --- | --- | --- | --- |
| | | | enol → ketone | ketone → enol | |
| Tris-HCl (2 mM) | 8.1 | 0.80 | 0.27 | 0.28 | - |
| Tris-HCl (10 mM) |  | 0.80 | 0.35 | 0.37 | 20/21 |
| Tris-HCl (30 mM) |  | 0.80 | 1.07 | 1.05 | 36/35 |
| Tris-HCl (60 mM) |  | 0.80 | 1.98 | 2.00 | 37/37 |
| Tris-HCl (100 mM) |  | 0.80 | 2.8 | 2.9 | 33/34 |
| HEPES-KOH (10 mM) | 7.5 | 0.50 | 2.3 | 2.5 | 420/460 |
| HEPES-KOH (100 mM) |  | 0.50 | >10 | >10 | >200 |
| Na-phosphate (10 mM) | 7.2 | 0.33 | 0.47 | 0.51 | 82/94 |
| MOPS-KOH (10 mM) | 7.2 | 0.33 | 1.1 | 1.1 | 230/230 |
| Imidazole-HCl (10 mM) | 7.0 | 0.24 | 0.74 | 0.74 | 310/310 |
| PIPES-KOH (10 mM) | 6.8 | 0.15 | 0.77 | 0.70 | 500/450 |

<sup>a</sup> $pK_a$ value of the conjugate acid.
<sup>b</sup> Fraction of the conjugate acid in the total buffer concentration at pH 7.5.
<sup>c</sup> The errors did not exceed 10%.
<sup>d</sup> Second-order rate constant for the reaction catalyzed by the acidic component of the buffer. Values separated by the slash were derived from the $k_{obs}$ values measured for the enol-to-ketone/ketone-to-enol conversions.

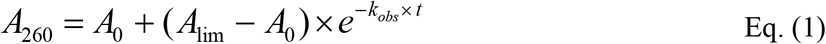

The *k*_obs_ value thus obtained represents the sum of the constants *k*^+1^ and *k*^−1^ (Fig. 1); however, since the tautomerization equilibrium in aqueous solution is strongly shifted toward the keto form (*k*^+1^ >> *k*^−1^), *k*_obs_ did not differ appreciably from *k*^+1^. The *k*_obs_ values measured in this way in several common buffers for the tautomerization reaction proceeding in the opposite directions (Table 1) were the same within the error of determination.

pH buffers are known to accelerate oxaloacetate tautomerization [26], and our data are fully consistent with this notion (Table 1). In the absence of a buffer, the *k*_obs_ value was low but not zero, indicating that the reaction also proceeds, but very slowly. Extrapolation of the dependence of *k*_obs_ on Tris buffer concentration to zero yielded a value of 0.20 ± 0.05 min^−1^ for the reaction catalyzed by water alone (Fig. 2D). This value is substantially lower than in the presence of 10 or 100 mM any of the buffers tested (Table 1). An unexpected finding was the anomalously high efficiency of HEPES as a tautomerization catalyst at pH 7.5 compared to Tris and other buffers. The observed rate constant *k*_obs_ in the presence of HEPES was almost an order of magnitude greater than in the presence of equal concentrations of Tris, previously used routinely for measuring OAD activity [9, 23, 24].

According to the Brönsted equation, the efficiency of general acid-base catalysis correlates with the p*K*_a_ value of the catalyst [28, 29]. The Brönsted plot constructed from the data in Table 1, taking into account the degree of buffer protonation (Fig. 2D), had a negative slope, indicating general acid catalysis (transfer of a proton to the carbon atom of oxaloacetate from the protonated form of the buffer) [29]. The slope of the line, which can vary from 0 to 1, was 0.53 ± 0.03, which is typical for such reactions [30].

The point corresponding to HEPES deviated strongly from the line obtained for the other buffers. Such deviations are not uncommon in Brönsted plots and may have several explanations, as discussed in Jencks’ monograph [30]. It is also possible that the exceptional efficiency of HEPES is explained by its zwitterionic nature, which, together with the structural features of its molecule, allows it to simultaneously perform acid and base catalysis, as required for keto-enol tautomerization [31]. Other zwitterionic buffers, MOPS and PIPES, if they possess such an ability, do so to a lesser extent (Table 1 and Fig. 2D). A contribution of the protonated sulfonic group to general acid catalysis cannot be ruled out either.

Importantly, at 100 mM HEPES, the tautomerization of oxaloacetate proved to be sufficiently fast (Fig. 2A and 2B) to preclude partial rate limitation of the enzymatic reaction even at high OAD concentrations in the assay medium. Therefore, all subsequent measurements of OAD activity were carried out in the presence of 100 mM HEPES.

### 3.2. Revision of steady-state kinetic data of OAD

Using the assay conditions optimized above, the kinetic parameters of the OAD-catalyzed oxaloacetate decarboxylation reaction were measured. Previously, OAD-2 activity in membrane preparations was observed only in the presence of Na^+^ ions [9], and we confirmed this result.

According to the same report, the dependence of *V. cholerae* OAD-2 activity on Na^+^ concentration is described by a sigmoidal curve with Hill coefficient and *K*0.5 values of 1.8 and 1 mM, respectively [9]. According to our data (Fig. 3), the Hill coefficient is 1.0, i.e., Na^+^ binding is non-cooperative, and the dependence is well described by the Michaelis-Menten equation with a Michaelis constant *K*_m,app_ = 1.1 ± 0.1 mM. The reason for the discrepancy in the data is unclear. The absence of cooperativity appears to be important for understanding the transport mechanism, given the accepted transport stoichiometry (2 Na^+^ per substrate molecule [14]).

**Fig. 3.**
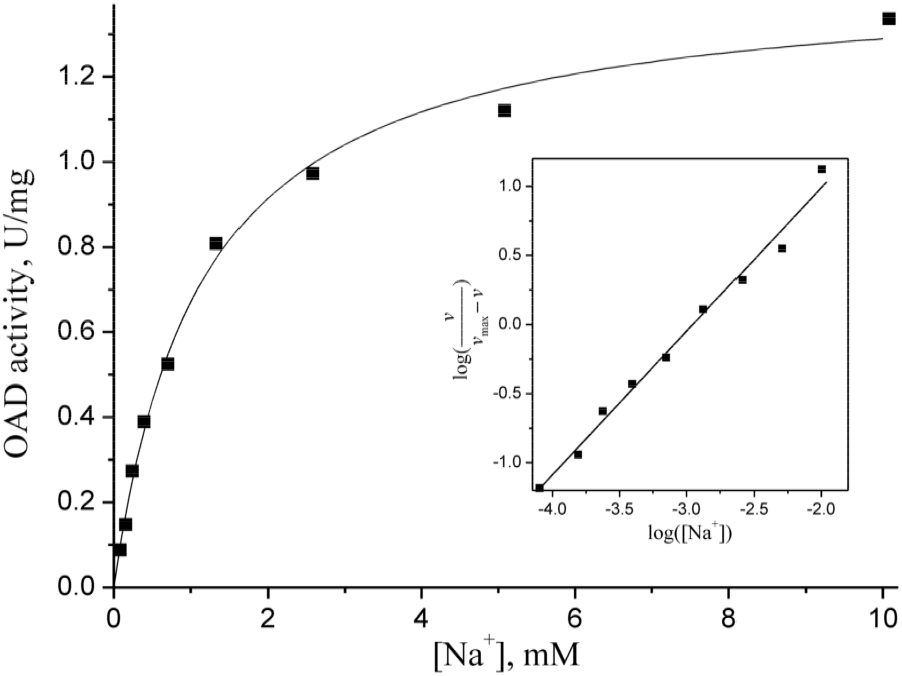
Dependence of OAD-2 enzymatic activity on Na^+^ concentration. Activity was measured spectrophotometrically by following the absorbance of oxaloacetate at 260 nm relative to 290 nm. Measurements were carried out in a buffer containing 100 mM HEPES-Tris (pH 7.5), 250 μM oxaloacetate, 50 μg/mL *V. cholerae* membrane vesicles, and varying concentrations of NaCl. The background Na^+^ concentration (prior to NaCl addition) was 80 μM. The oxaloacetate stock solution at pH 7.5 was prepared in water. The curve corresponds to the best fit of the Michaelis-Menten equation. The inset shows the same data plotted as a Hill plot, which yielded the Hill coefficient of 1.03 ± 0.04.

The dependence of the OAD-2 reaction rate, determined as for Fig. 3, on oxaloacetate concentration at a saturating Na^+^ concentration (20 mM) was hyperbolic (data not shown), as was previously reported for the *Klebsiella aerogenes* enzyme [32]. The *K*_m_ value for oxaloacetate was 56 ± 1 μM in terms of total oxaloacetate concentration, which is close to the value previously obtained for OAD from *K. aerogenes* (90 μM). We also confirmed the competitive inhibition of OAD by the substrate analogue – oxalate [9, 32]. According to our data, oxalate reduced the apparent affinity of OAD-2 for oxaloacetate (Fig. 4) with a *K*_i_ value of 2.0 ± 0.2 μM. This value is close to the measured dissociation constant of 1.3 μM for the equilibrium binding of oxalate to OAD from *K. aerogenes* [32]. The *V*_max_ value was not affected by oxalate.

**Fig. 4.**
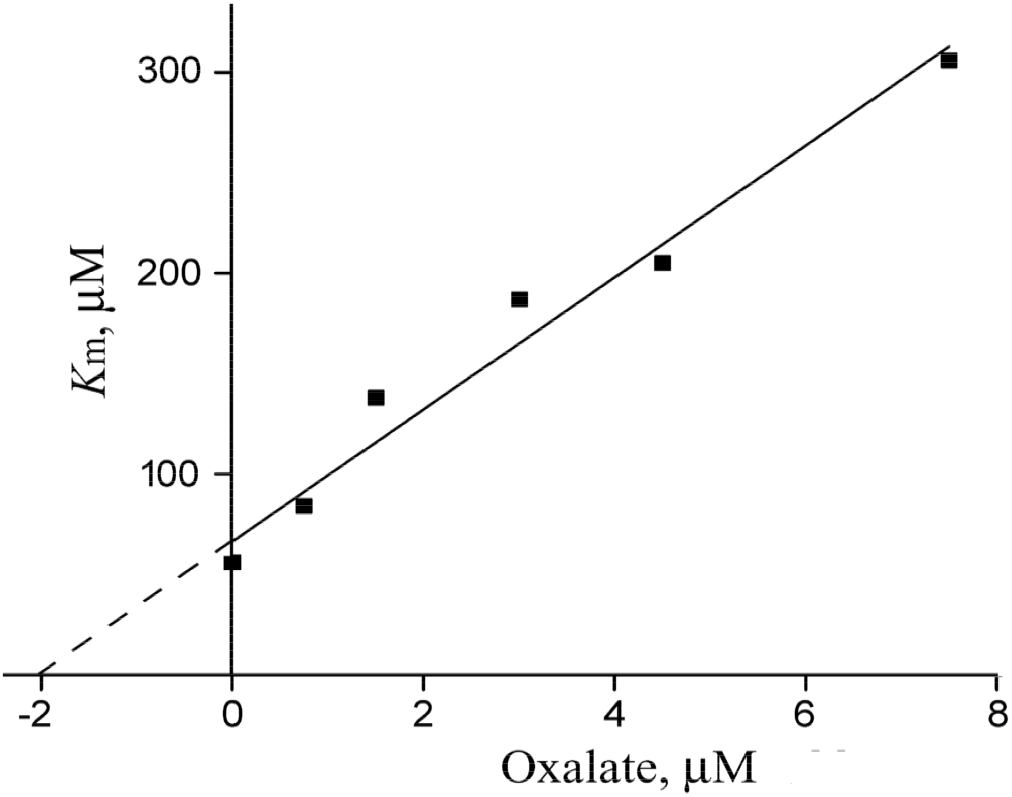
Dependence of the OAD-2 *K*_m_value for oxaloacetate on oxalate concentration.

### 3.3. Identification of the tautomeric form of oxaloacetate converted by OAD-2

Oxaloacetate exists in solution in two tautomeric forms—ketone and enol (Fig. 1). The available indirect evidence as to which form is the true substrate of OAD are contradictory. Thus, isotope exchange data point to the keto form [23]. In contrast, the kinetics of the enzymatic reaction under conditions where substrate tautomerization becomes a partially rate-limiting step at high enzyme concentrations can be interpreted as indicating that it is the enol form [15]. We addressed this controversy in a direct experiment using a coupled assay with LDH to measure OAD-2 activity. The assay monitored product (pyruvate) formation and, because the product is common for both tautomeric forms of the substrate, obviated the need to monitor the concentrations of both forms.

The OAD-2 used in these experiments was purified from the *V. cholerae* membrane fraction by means of affinity chromatography on avidin-agarose in order to remove interfering proteins. The resulting OAD preparation demonstrated a high Na^+^-dependent decarboxylase activity (62 μmol min^−1^ mg^−1^) and little contamination by polypeptides other than OAD-2 subunits on SDS-PAGE (Fig. 5A). The electrophoretic separation was performed without boiling the samples to avoid loss of hydrophobic subunits [33], which was particularly significant with OAD [24]. According to the data obtained (Fig. 5A), OAD-2 dissociation was incomplete and, along with the bands of individual α, β, and γ subunits, a band of the α_n_ subunit multimer was also present. MALDI-MS analysis confirmed the assignment of the bands to α, β, and γ subunits of OAD-2 and α_n_ multimer (sequence coverage of 67, 30, 40, and 69%, respectively) (KEGG IDs: VC0395_A0319–VC0395_A0321).

**Fig. 5.**
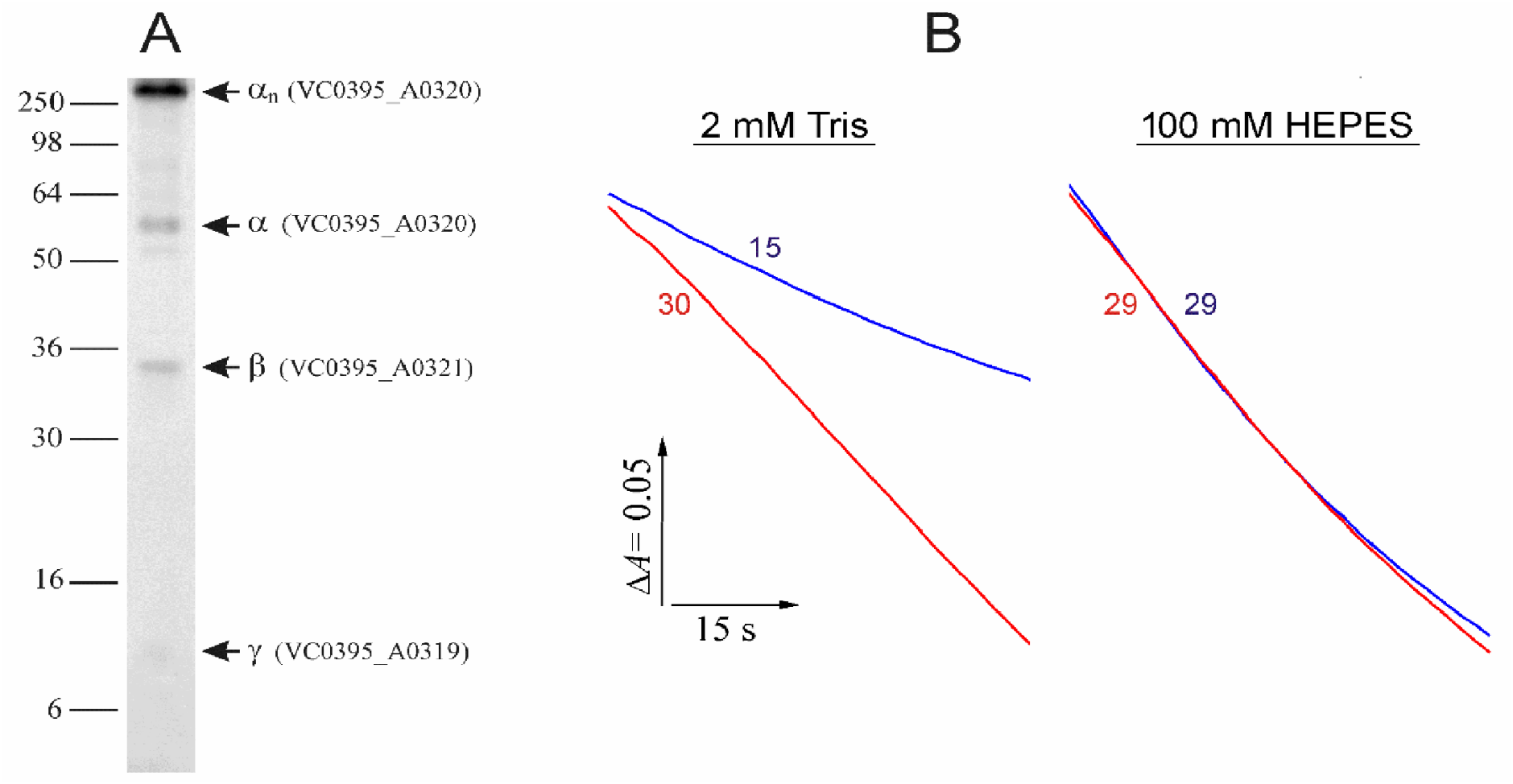
Identification of the converted form of oxaloacetate. **(A)** Characterization of the purified OAD-2 preparation by SDS-PAGE. Protein load was 6 μg per lane. The bars with numbers on the left side indicate the positions and molecular masses of marker proteins. The OAD-2 subunits identified by MALDI-TOF-MS analysis are indicated on the right side. **(B)** Kinetic traces of oxaloacetate conversion by the purified OAD-2 preparation. The assay medium contained 20 mM NaCl, 1 U/mL LDH, 0.1 mM NADH, and either 2 mM Tris-HCl or 100 mM HEPES-KOH, as indicated above the traces. The reaction was initiated by the addition of 50 μM oxaloacetic acid from a stock solution in water (red traces) or in dry diethyl ether (blue traces). The resulting pH value was 7.5 in both cases. The reaction was monitored by absorbance at 340 nm (maximum in the NADH spectrum). OAD-2 activities in U/mg are indicated next to the traces.

The identification of the tautomeric form of oxaloacetate that is the true substrate of OAD-2 was possible due to two reasons: the ability to vary the relative content of the tautomeric forms in the substrate stock solution (by changing the solvent and pH) and the low rate of tautomerization in the reaction medium in the presence of 2 mM Tris-HCl buffer, pH 7.5 (Fig. 2). Under these conditions, we compared the rates of conversion of 50 μM oxaloacetic acid added to the reaction medium from the stock solution in diethyl ether (67% enol and 33% ketone forms) and acidic aqueous solution (6.5% enol and 93.5% ketone forms).

Switching from a stock solution in diethyl ether to a stock solution in water, while keeping the total substrate concentration constant, increases the initial concentration of its keto form and decreases the initial concentration of the enol form. According to the Michaelis-Menten equation, the rate of the enzymatic reaction at a non-saturating substrate concentration should increase if the true substrate is the keto form of oxaloacetate, and decrease if the enol form is the true substrate. As shown in Fig. 5B (left panel), the reaction rate increased approximately twofold, indicating the conversion of the keto form of the substrate. Such a preference is also characteristic of many other enzymes that use oxaloacetate as a substrate [34].

A simple calculation shows that if the keto form is the true substrate and the enol form is completely inert (neither binds nor is converted), the ratio of the rates obtained with the indicated substrate stock solutions should be 1:1.94. Indeed, the Michaelis constant determined above (56 μM) refers to the sum of the two tautomeric forms of the substrate. If only the keto form is converted, which in these experiments constituted 100% – 13% = 87%, then the true *K*_m_ value for it is 56 μM × 0.87 = 48.7 μM. The concentration of the keto form in the reaction medium in the experiments with the ethereal stock solution of the substrate was 50 μM × 0.33 = 16.5 μM, and in the experiment with the aqueous stock solution of the substrate — 50 μM × 0.935 = 46.8 μM. According to the Michaelis-Menten equation, the “water/ether” reaction rate ratio would be (1 + 48.7/16.5) / (1 + 48.7/46.8) = 1.94. The experimentally determined ratio is 2.0 (Fig. 5B, left panel), i.e., it agrees with the theoretical value within experimental errors. This result confirms that only the keto form of the substrate is converted. Moreover, this result indicates that the enol form is inert, i.e., it is not a competitive inhibitor, since otherwise the measured ratio of the two rates would be significantly greater than that calculated theoretically due to the higher content of the enol form in the ethereal solution. At a minimum, these data indicate that the inhibition constant for the enol form of oxaloacetate exceeds 0.5 mM.

When the reaction medium contained a 100 mM HEPES buffer that ensured a high rate of keto-enol tautomerization (Fig. 2), the rates of oxaloacetate conversion were identical (Fig. 5B, right panel), i.e., they did not depend on the ratio of the keto and enol forms in the substrate stock solution. We also note that OAD-2 did not accelerate the keto-enol tautomerization of oxaloacetate in a medium with a low buffer concentration (2 mM Tris-HCl, pH 7.5) lacking Na^+^ ions required for the decarboxylation reaction.

### 3.4. The source of the proton in the catalytic cycle of OAD-2

The decarboxylation of oxaloacetate consumes a proton from the medium (Fig. 1), making the proton a participant in this chemical reaction [11]. Di Berardino and Dimroth [16] proposed a “direct”-coupling mechanism, which assumes that the “chemical” proton required for decarboxylation is taken up from the periplasmic side of the membrane, implying that OAD catalyzes an electrogenic 2Na^+^/1H^+^ exchange. This hypothesis was based on the observation of a valinomycin-resistant ΔpH generation (alkalization at the “periplasmic” side of the membrane) during OAD operation in proteoliposomes.

However, the alkalization of the internal volume of proteoliposomes during OAD operation can also be explained by electrophoretic transport of protons from the vesicle interior, driven by the transmembrane electric potential difference Δψ generated during Na^+^ transport [35]. Such secondary proton transport is well-documented in sodium pumps [36-39]. The two mechanisms of alkalization of the vesicle interior can be distinguished based on the following considerations [35, 40]. The transmembrane uptake of “chemical” protons must be sensitive to protonophorous uncouplers and insensitive to substances that reduce Δψ (e.g., K^+^ or Na^+^ ionophores) or even be stimulated by them. In contrast, the electrophoretic transport of protons will be sensitive to K^+^-or Na^+^-ionophores but insensitive to protonophorous uncouplers, or may even be stimulated by them. Di Berardino and Dimroth [16] found that the alkalization observed during OAD operation is only weakly sensitive to valinomycin, which precludes a reliable distinction between the alternative mechanisms.

We performed similar measurements using *V. cholerae* membrane vesicles containing entrapped pyranine, a fluorescent pH indicator which directly senses proton accumulation in or efflux from the vesicles. Addition of oxaloacetate to the vesicles led to a rapid alkalization of their internal volume (Fig. 6A, blue trace). This effect was not observed in the absence of Na^+^ ions (Fig. 6A, purple trace), suggesting the involvement of OAD-2 activity in ΔpH formation. The presence of the protonophore CCCP in the assay medium led to a slight stimulation of the oxaloacetate effect (Fig. 6A, red trace), indicating a major role for electrophoretic proton transport in the measured signal. Consistent with this conclusion, the addition of the K^+^-ionophore valinomycin suppressed the signal (Fig. 6A, black trace); however, this suppression was only partial, consistent with the findings of Di Berardino and Dimroth [16].

**Fig. 6.**
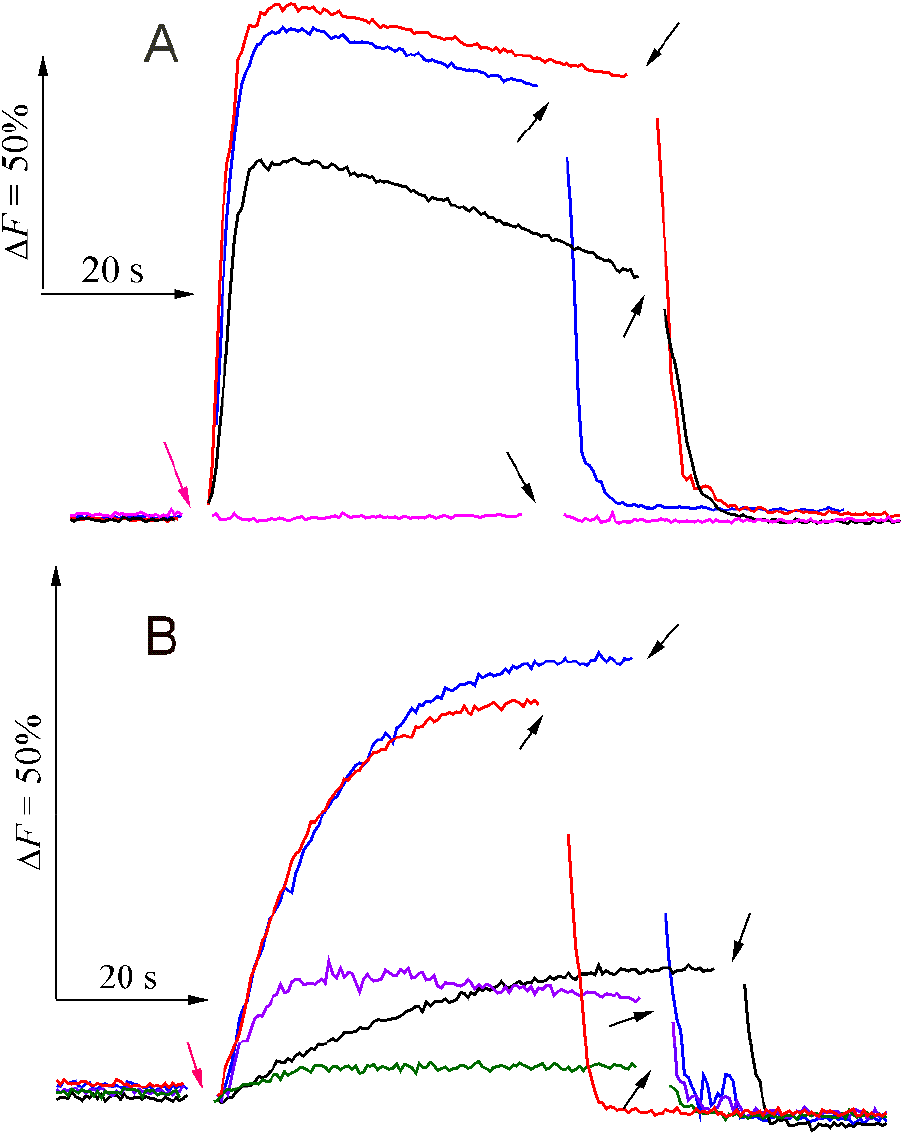
Fluorescence of entrapped pyranine in *V. cholerae* OAD-2-containing membrane vesicles. An increase in fluorescence indicates alkalization of the internal volume of the vesicles. **(A)** In the absence of oxalate. The assay medium contained 100 mM MOPS-Tris (pH 6.5), 25 mM K_2_SO_4_, 5 mM MgSO_4_, 10 mM Na_2_SO_4_ (except for the purple trace), and membrane vesicles with entrapped pyranine (75 μg protein per mL). The background Na^+^ concentration (prior to Na_2_SO_4_ addition) was 10 μM. The reaction was initiated by the addition of 300 μM oxaloacetate (indicated by purple arrows). To dissipate ΔpH, 2 μM monensin (2 μM nigericin for the purple trace) was added at the time points indicated by black arrows. Blue trace, no additives; red trace, in the presence of 10 μM CCCP; black trace, in the presence of 1 μM valinomycin; purple trace, with no Na^+^ added. **(B)** In the presence of 100 μM oxalate. Violet trace, in the presence of 20 μM Na^+^-ionophore ETH157; green trace, in the presence of 1 μM valinomycin and 20 μM ETH157. Other conditions and labels are as in panel A.

To investigate this further, we hypothesized that the low sensitivity of the H^+^ transport to valinomycin resulted from rapid depletion of potassium ions inside the vesicles. Based on the vesicle volume of ∼0.5 μL/mg protein and OAD-2 activity of 1.3 µmol/min per 1 mg protein, one can estimate that the K^+^ concentration inside the vesicles would decrease via exchange for Na^+^ at a rate of approximately 40 mM/s. This value is very high, but it represents only an upper limit because this calculation did not account for the possibility of Na^+^ and other ion leakage from the vesicles.

To test this hypothesis, we repeated the experiment in Fig. 6A in the presence of 100 μM oxalate, a competitive inhibitor of OAD-2, which, at an oxaloacetate concentration of 300 μM, reduces the enzyme activity approximately 10-fold (see section 3.2.). As a result, the oxaloacetate-driven alkalization was reduced, as expected, both in the amplitude and the rate of ΔpH build-up (Fig. 6B, blue trace) but remained insensitive to CCCP (Fig. 6B, red trace). Importantly, the effect was substantially suppressed in the presence of a K^+^- or Na^+^-ionophore (Fig. 6B, black and violet traces) and was nearly completely suppressed when both ionophores were present (Fig. 6B, green trace). Notably, neither valinomycin nor ETH157 inhibited OAD-2 activity at the concentrations used.

Our findings thus provided strong evidence that the partial effect of valinomycin is indeed explained by K^+^ depletion and that the ΔpH formation associated with OAD-2 operation is caused by the electrophoretic transport of protons. Hence, the “chemical” proton required for decarboxylation is taken up by the enzyme from the cytoplasmic side of the membrane.

### 3.5. Conclusions

We propose an improved method for measuring OAD (oxaloacetate decarboxylase) activity. The improvement is based on the newly identified ability of HEPES buffer to act as an exceptionally efficient general acid catalyst of the keto-enol tautomerization of oxaloacetate. The proposed method may also be useful for measuring the activity of other oxaloacetate-consuming (or-producing) enzymes. Using this approach, we established that OAD decarboxylates exclusively the keto form of oxaloacetate and in this respect resembles most other oxaloacetate-converting enzymes.

Furthermore, our results demonstrate that the ΔpH generated by OAD is formed exclusively through secondary electrophoretic transmembrane proton transport. A clear implication is that the “chemical” proton consumed during the decarboxylation reaction is taken up from the cytoplasm.

This finding advances our knowledge of the mechanism of OAD action and rules out the “direct”-coupling model previously proposed for this enzyme [16]. Thermodynamically, the established proton pathway would allow more energy from the OAD-catalyzed decarboxylation reaction to be conserved.

Based on these findings, the equation for the reaction catalyzed by OAD can be revised as:

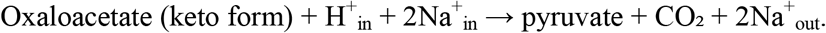

## CRediT authorship contribution statement

Yulia Bertsova: Investigation, Visualization. Alexander Kvartalov: Investigation. Marina Serebryakova: Investigation. Alexander Baykov: Data curation, Visualization, Writing. Alexander Bogachev: Conceptualization, Methodology, Investigation, Writing.

## Acknowledgements

MALDI MS facility became available to us in the framework of the Moscow State University Development Program PNG 5.13. This work is dedicated to the memory of Prof. A.D. Vinogradov and his contribution to the study of keto-enol tautomerization.

## Data availability

All data are available from the authors at a reasonable request.

